# High-resolution spatial transcriptomics of stem and storage root vascular cambia highlights key regulatory processes for xylem parenchyma differentiation in cassava

**DOI:** 10.1101/2025.01.24.634672

**Authors:** David Rüscher, Uwe Sonnewald, Wolfgang Zierer

## Abstract

Due to their high carbohydrate content, the storage roots of cassava are an important food source for hundreds of millions of people worldwide. In contrast to the woody stems of the plant, the xylem of the storage roots produces mainly starch-rich storage parenchyma cells and only few tracheary elements and almost no fibers. Despite these obvious differences, both stems and storage roots are formed by a vascular cambium. To find more insights into the differences in the regulation of cell division and differentiation in stems and storage roots, a cryo-sectioning approach was utilized, to generate high-resolution transcriptome profiles spanning the entire vascular cambium of both tissues. We observed that storage parenchyma formation is connected to the repression of secondary cell wall formation through a decrease in expression levels of key players in the NAC/MYB regulatory network, as well as decreases in the downstream pathways for lignin and hemicellulose biosynthesis. Additionally, the expression of *MeWOX14*, a transcription factor associated with GA signaling and xylem fiber differentiation, is strongly reduced in storage roots compared to stem xylem. By contrast, the expression of *MeKNOX1*, a well-known meristem regulator, as well as most cassava *LSH* genes and several ABA-related transcription factors were associated with parenchyma cells. Our data suggest that the repression of secondary cell wall formation and GA signaling, together with an active auxin and ABA signaling, as well as extended *MeKNOX1* activity could control storage parenchyma formation in cassava storage roots.

**Significance statement:** Tissue-specific expression data is still scarce for cassava and the regulatory mechanisms controlling the formation and differentation of secondary vasculature cells are largely unknown in this species. By performing a cryosectioning approach on cassava stems and storage roots, we generated highly-resolved, tissue-specific transcriptomic data, identified key factors for parenchyma cell formation in storage roots and propose a working model for further research.

## Introduction

Cassava (*Manihot esculenta*) is a staple food crop whose storage roots represent an important food source for hundreds of millions of people in Sub-Saharan Africa, South America, and Southeast Asia. Despite several advances in the understanding of the plant’s physiology and biochemistry, the regulatory mechanisms governing the development of the cassava storage roots are still not well understood, although this knowledge might greatly assist the breeding of specific root traits or biotechnological approaches targeting storage root quality and yield.

Cassava storage roots are initiated from stem-derived roots that eventually initiate secondary growth (Chaweewan and Taylor, 2015; Mehdi *et al*., 2019; Rüscher *et al*., 2021). Their cellular anatomy is overall similar to their stem or storage root neck tissues, but they possess much less xylem vessels and mostly xylem parenchyma cells instead of xylem fibres. Several bulk tissue studies have already described the storage root initiation and root bulking process (e.g. Chaweewan and Taylor (2015); Rüscher *et al*. (2021); Rüscher *et al*. (2024); Siebers *et al*. (2017); Sojikul *et al*. (2015)), but a clear understanding of the regulatory processes leading to the differentiation of starch-storing parenchyma cells over xylem fiber cells in storage roots is still missing. While single cell transcriptomic analysis would be best suited, this approach faces technical challenges. The presence of starch granules in parenchyma cells will lead to a rapid rupture or isolated protoplasts, leading to a strong negative selection against storage parenchyma cells. Moreover, single cell transcriptome workflows rely on cell type-specific marker genes that are currently not well-established in cassava.

To gain more insight into tissue-specific gene expression in cassava, we performed a comparative cryo-sectioning approach of cassava stem and storage root tissues to identify commonalities and differences in the regulation of cell division and differentiation in these two organs. While this method is difficult on irregularly-shaped organs, the symmetrical nature of cassava storage roots makes it well-suited. Cryosectioning approaches were successfully used on tree stems, where these were mounted on a cryo-microtome to produce sequential sections, which were either used for RNA-sequencing or microarray analysis (Immanen *et al*., 2016; Schrader *et al*., 2004; Sundell *et al*., 2017). Consequently, the expression profile of the most important regulators could be studied in detail, leading to an improved understanding of wood formation. By performing a comparative cryo-sectioning analysis of cassava stem and storage root tissues, this study presents additional insights into xylem parenchyma formation in cassava storage roots.

## Results

### Generation of meristematic, transition zone, and fully-differentiated tissue sections in stems and storage roots

The vascular cambium area of cassava stems and storage roots consists of multiple specialized zones. The actively dividing cambial zone contains only a few cell layers and is approximately only 80 µm thin (Fig. 1). It encompasses ray initials, producing vascular rays, and fusiform initials, producing phloem and xylem mother cells. In both stem and storage root tissues, phloem mother cells undergo differentiation almost immediately. In stems, xylem mother cells start differentiating post-division, initiating secondary cell wall formation in the xylem transition zone. There, forming tracheary elements rapidly enlarged, while fibers and parenchyma mostly maintain their size. Eventually, programmed cell death of tracheary elements and fibers is initiated, and their cell walls rapidly grow and lignify. A sharp border lies between the differentiated xylem and the xylem transition zone of the stem, which can be made visible by toluidine blue staining (Fig. 1). The xylem transition zone of the storage root looks much different in comparison and is much harder to identify, as there is no clear-cut border between newly generated and fully-differentiated xylem cells. Except for a few tracheary elements the cells mainly undergo growth and rapidly initiate starch production forming the so-called storage parenchyma (Fig. 1). These clear difference in anatomy and development of stem and storage root xylem already highlights differences in the underlying genetic regulation. To decipher the transcriptional differences between stem and storage roots, a high-resolution RNA-seq dataset was produced using 20 µm thin sequential samples produced through cryosectioning, as described in Schrader *et al*. (2004). Up to 17 sections of three individual stems and storage roots were gathered, spanning from −100 µm (outwards) to 220 µm (inwards) from the cambium in the center (Fig. 1).

**Figure 1.**
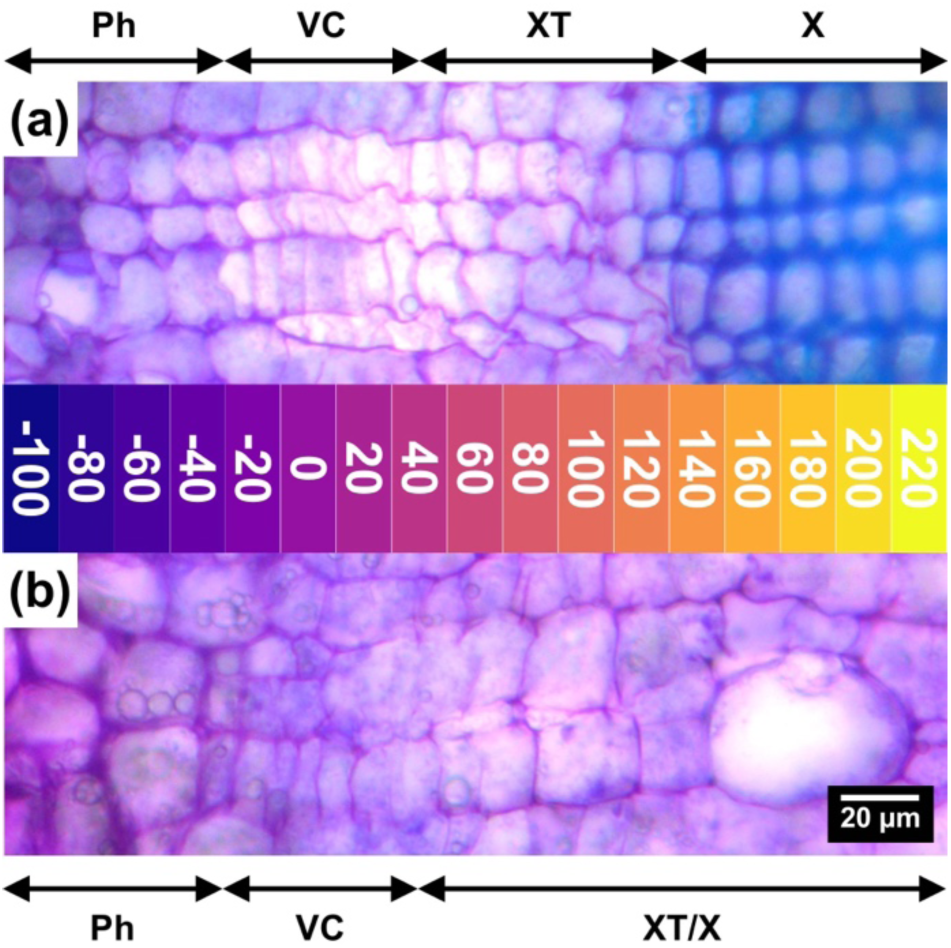
Overview of the cryosectioning approach. Overview of the cryosectioning approach of stems (a) and storage roots (b). Each colored block indicates a 20 µm cryomicrotome slice used for later analysis. Numbers indicate the distance from the cambium in µm. Arrows depict the different tissues. Micrographs are toluidine-blue stained hand-sections before cryosectioning. Ph, phloem; VC, vascular cambium; XT, xylem transition zone; X, xylem.

These sections comprised the actively dividing area of stem and storage root tissues, as well as the xylem transition zone and differentiated phloem and xylem cells. The generated RNA-seq data showed a clear trend in the sections from phloem to xylem visualized using a low-dimensional representation (Fig. 2a). This trend, however, was divergent for the different plants. This was exacerbated for storage roots, likely due to their slightly convex shape compared to the straight stems. Therefore, a linear regression model was fitted to the data, with the batch effects of the individual plant subtracted, while maintaining the information about the distance from the cambium. This mitigated the systematic differences between samples. However, simply using the distance as a design variable would not yield much useful information, due to a visible shift in the gradient from phloem to xylem sections between specimens (Fig. 2b). Since the objective is not to assess changes relative to distance per se (which are slightly different for the individual stems analyzed), but rather to identify expression profiles across the various zones of the vascular cambium, clustering of samples was performed to identify groups of samples with similar expression profiles using community detection (Blondel *et al*., 2008) on a *k*-nearest neighbor graph. This methodology is similar to finding cell clusters in single-cell transcriptomics (Fig. 2c; see Methods for detail). Four clusters were identified for each tissue, named A to D, with increasing median distances per cluster. These clusters exhibited a clear linear increase in distance from the cambium for each tissue (Fig. 2d). Consequently, these clusters should capture the required information about the distinct zones of the cambium. Therefore, the clusters will be referred to as “phloem”, “phloem/cambium”, “cambium/xylem” and “xylem” for easier readability (Fig. 2d).

**Figure 2.**
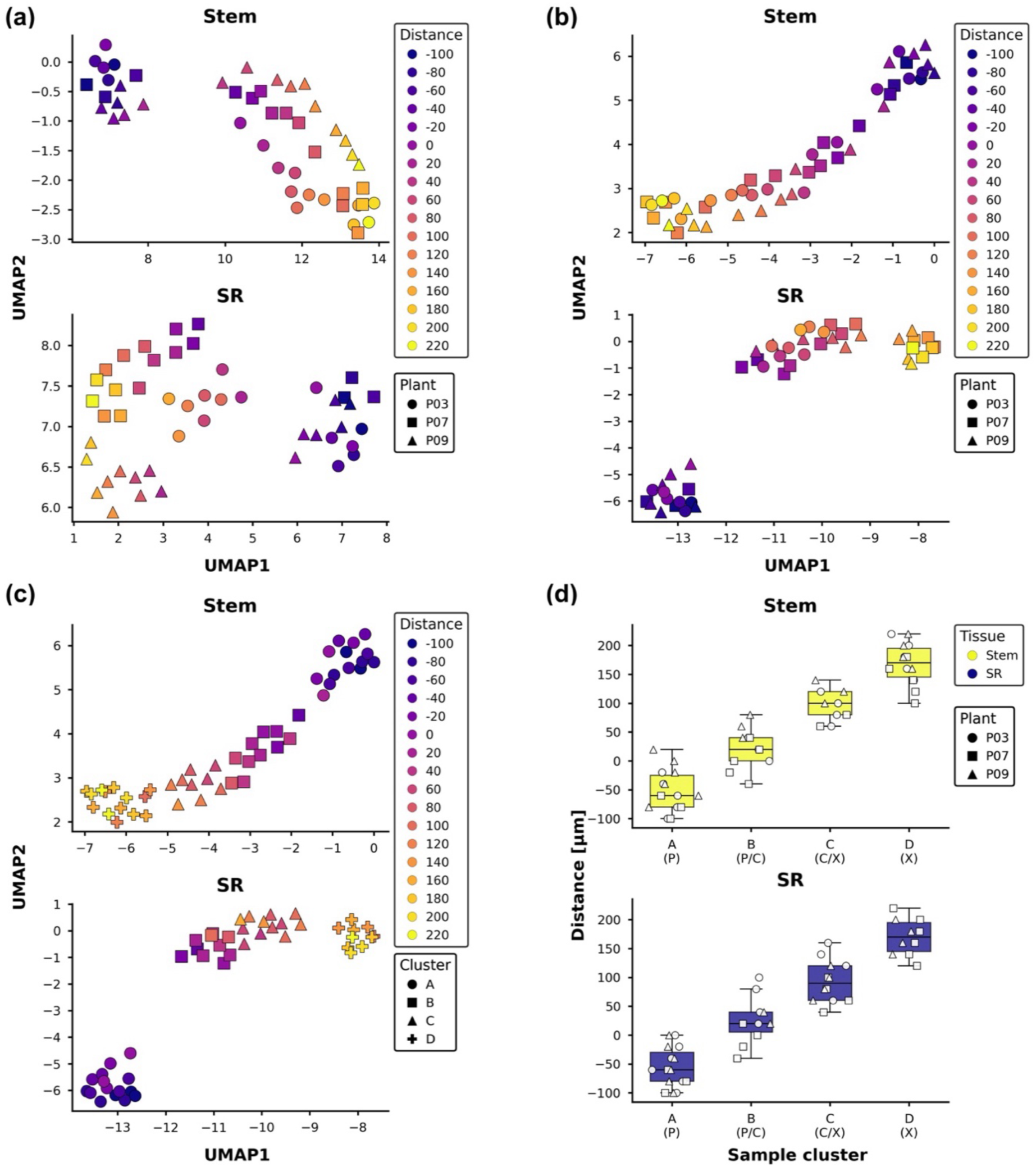
Section clustering approach. Overview of the clustering approach. 2D visualization of samples based on all expressed transcripts (normalized counts >50) using a dimensional reduction projection (UMAP; 45 neighbors, 0.001 minimum distance, and cosine distance) on (a) VST counts before processing and (b, c) after correcting for the systematic differences between plants. Shapes indicate (a, b) the plant and (c) the cluster. (a-c) Colors indicate the distance from the cambium. (d) Boxplot depicting the distance from the center of the cambium for each cluster. Different shapes refer to the different plants. Abbreviations: P: phloem; C: cambium; X: xylem; SR: storage root.

The newly identified phloem, phloem/cambium, cambium/xylem and xylem clusters (Fig. 2c-d) were used as design variable in a likelihood ratio test (LRT) to find differentially expressed genes (DEGs) within each tissue. A total of 18,032 and 15,381 DEGs (adjusted p-value < 0.001) were identified for stem and storage root, respectively (Supplementary File 1). Since the LRT does not give information about the actual behavior of the DEGs across the sample clusters, a clustering approach was used to find groups of DEGs with similar expression profiles (Fig. 3a-b; Supplementary File 2). This was done by generating a co-regulation network based on Pearson correlation coefficients and community detection (see Methods for details). Five clusters with similar expression profiles were found in both tissues (Fig. 3a-b). Genes with higher expression in the phloem were found in clusters Stem 4 (6,114 DEGs) and SR 3 (4,273 DEGs), while Stem 3 and SR 1 expression increased towards the xylem. Therein, remarkably fewer DEGs were found in the storage root with only 2,224 compared to 4,743 in the stem. A similar number of genes was expressed in the respective cambium/xylem cluster with 4,400 in Stem 2 and 4,751 DEGs in SR 4 (Fig. 3c). Only few DEGs peaked towards the phloem/cambium cluster, with 1,238 in Stem 1 and 245 in SR 0. Comparing the contents of the expression clusters across stem and storage root tissues showed that overall, the intersections were largest between clusters with similar expression profiles, as expected (Fig. 3c). This was most visible in the overlap of the respective phloem and cambium/xylem expression clusters, which shared 2637and 2574 DEGs between stem and storage root tissues. However, clear differences in the overlap between stem and storage roots were also found. The stem phloem expression cluster for instance, also shared 1082 genes with the storage root phloem/xylem expression cluster (Fig. 3c).

**Figure 3.**
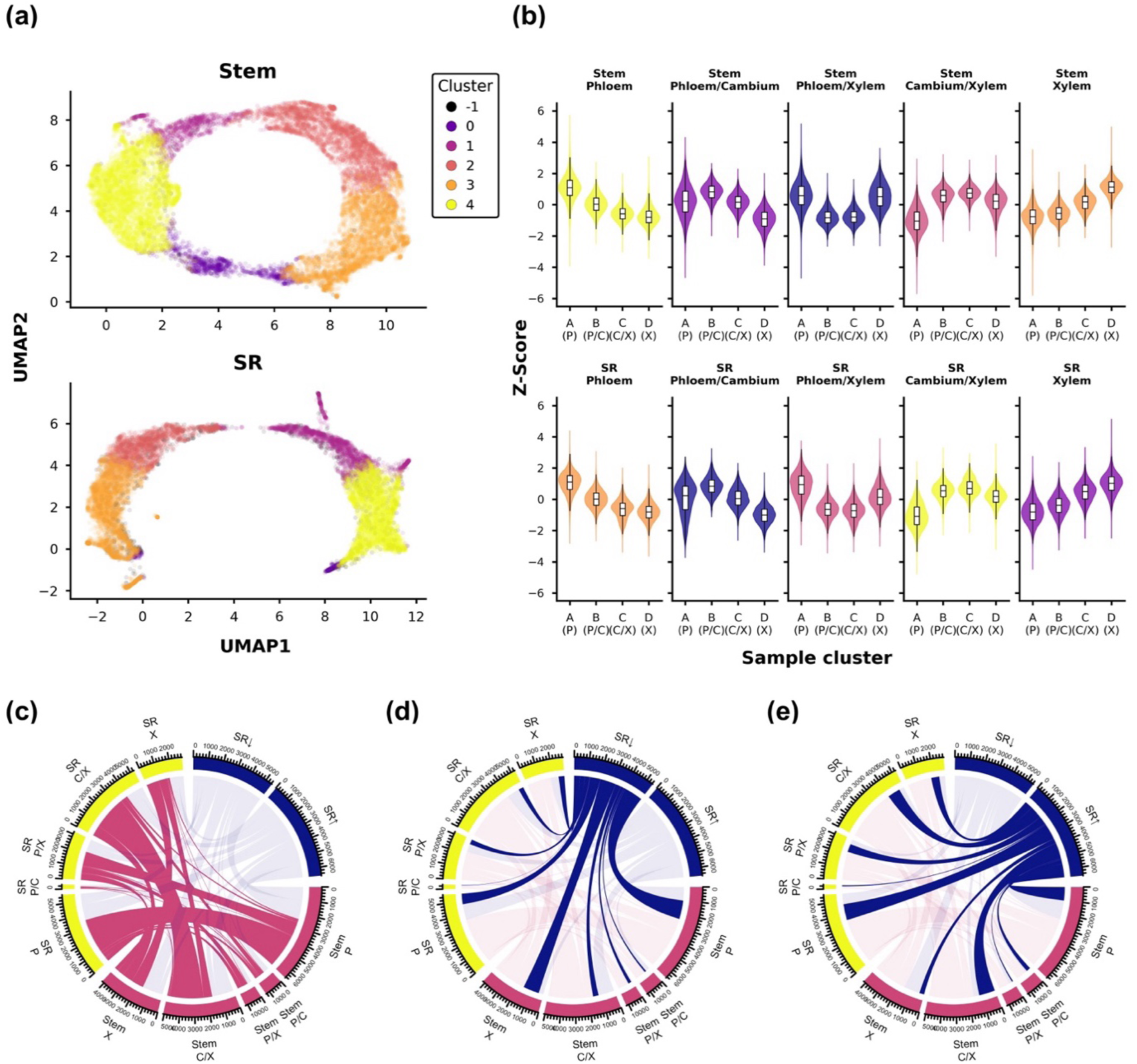
Clustering of gene expression. (a) Visualization of a co-regulation network analysis for DEGs (p-adjusted (FDR) < 0.001) within a tissue using a UMAP projection for (top) stem and (bottom) storage root sections. Each point represents a gene and colors indicate the gene cluster. Cluster −1 describes outliers. (b) Violin plots depicting the expression profiles for each cluster and tissue across sample clusters. Legend is shared with (a). (c-e) chord diagrams displaying the relationship between clusters. The thicker the line connecting two sets, the larger the size of the intersection between them. Storage root low and storage root high means genes are significantly lower or higher expressed in storage root compared to stem independent of the sample cluster. Highlighted were the connections from the same diagram between (c) stem and storage root gene clusters, (d) genes lower expressed in storage root and the gene clusters, and (e) genes higher expressed in storage root. Abbreviations: SR: storage root, SR↑: higher in SR, SR↓: lower in SR, P: phloem, C: cambium, X: xylem.

The clustering approach clearly showed that it is possible to find qualitative difference in gene expression between the two tissues, but some key regulations might come from quantitative differences of certain genes. Therefore, a Wald test was executed using tissue information as design variable, while controlling for the sample cluster to find DEGs between storage root and stem (Supplementary File 3). This resulted in 8,836 identified DEGs (adjusted p-value < 0.001 and |log2 fold-change| > 1.5), with 4,297 being higher expressed and 4,539 lower expressed in storage roots. Comparing DEGs from the Wald and LRT analysis revealed that 1,221 genes that are significantly lower expressed in storage roots were part of the stem xylem expression profile (Fig. 3d). Unsurprisingly, gene ontology (GO) term enrichment (Supplementary File 4) on these genes revealed an overrepresentation of biological processes involved in secondary cell wall biosynthesis. Conversely, genes with higher expression in storage roots (Fig. 3e) could mainly be found in the storage root cambium/xylem cluster (969 DEGs) and less frequently in the stem xylem cluster (365 DEGs). These showed an enrichment in primary cell wall biosynthesis and cell division (Supplementary File 5). Overall, these observations likely reflect the stronger growth of the storage root combined with the decrease in woody cell deposition and is consistent with earlier results obtained by comparing storage roots and woody fibrous roots (Rüscher *et al*., 2021; Rüscher *et al*., 2024).

### Marker gene expression confirms the location of clusters within the vascular cambium

Transcriptomics on sequential sections has the advantage that the rough position within the sample is known. Unlike in single-cell transcriptomics, this allows for correct identification of expression profiles without intricate downstream analysis (Fig. 3b). Nevertheless, the presence and expression profile of certain well-studied marker genes can still provide insight into the biological interpretability of the expression and sample clusters and can also reveal notable expression differences (Fig. 4).

**Figure 4.**
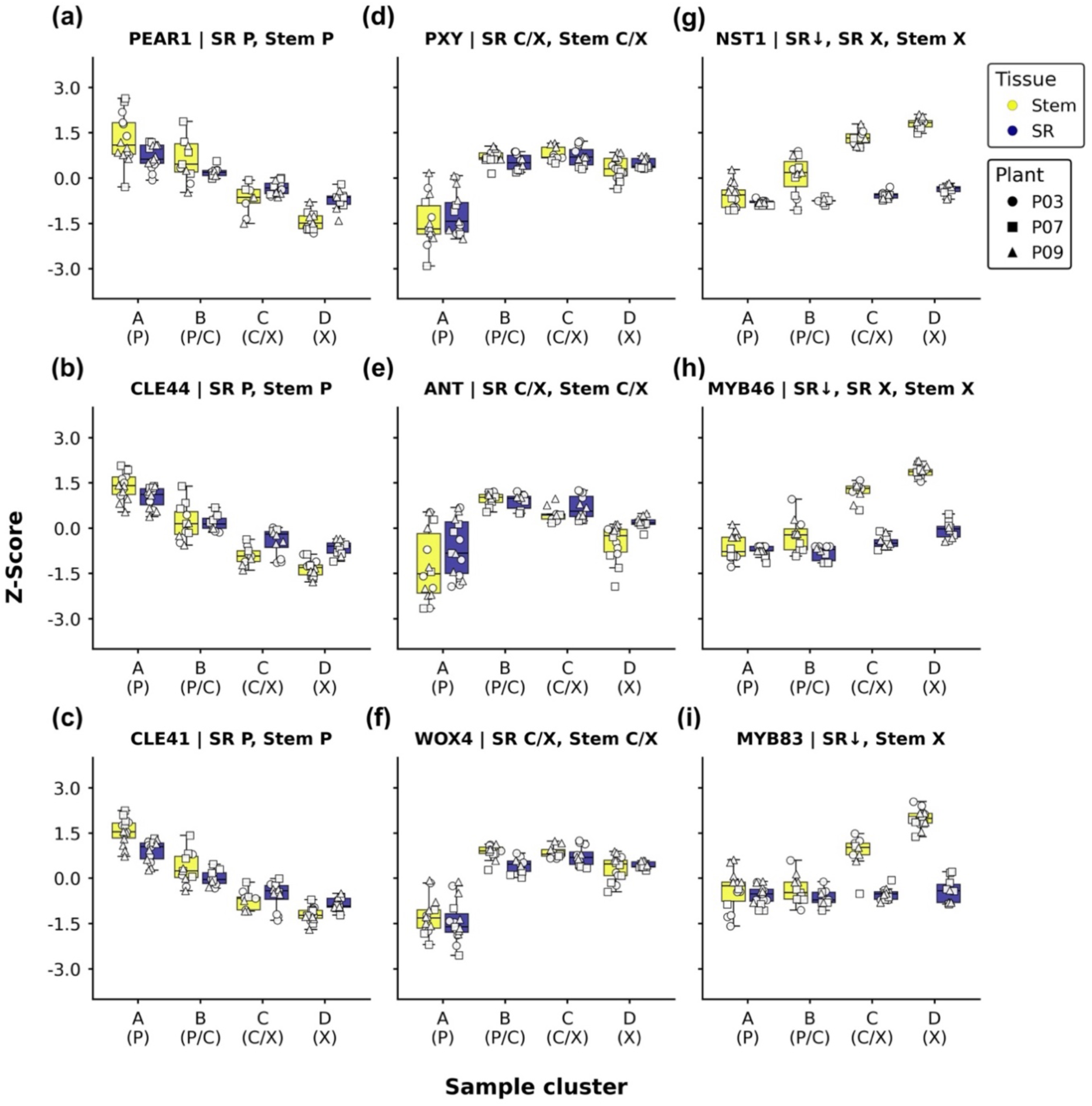
Expression of marker genes. Shown is the expression of well-studied marker genes against the sample cluster. Data is centered and scaled based on batch corrected VST values. The selected genes are reciprocal BLASTP hit against the *A. thaliana* Araport11 proteome. Columns from left to right show phloem, cambium, and xylem marker genes. Panel titles indicate the trivial name and cluster association. Abbreviations: SR: storage root, SR↑: higher in SR, SR↓: lower in SR, P: phloem, C: cambium, X: xylem.

Firstly, one would expect certain genes important for phloem development in the phloem expression clusters. For example, PHLOEM EARLY DOF (PEAR) are mobile transcription factors required for radial patterning and work antagonistic to class III HOMEODOMAIN LEUCINE ZIPPER (HD-ZIPIII) proteins in the differentiating phloem cells (Miyashima *et al*., 2019). Expression of the cassava *PEAR1* gene (Fig. 4a) was found to increase toward the phloem and could be grouped in the phloem clusters of both tissues and did not show significant differences between them. This was expected, considering the similarities in the phloem of stem and storage root. The same was found for the *CLAVATA/ESR-RELATED (CLE) 41* (Fig. 4b) and *CLE44* (Fig. 4c) orthologs. *CLE41/44* encode for small peptides also known as TRACHEARY ELEMENT DIFFERENTIATION INHIBITORY FACTOR (TDIF). These are expressed in the phloem and can be bound by the TDIF receptor PHLOEM/XYLEM INTERCALATED (PXY), which is required to establish the position of the vascular cambium (Etchells and Turner, 2010).

*MePXY* (Fig. 4d) was used as a cambium marker gene and showed the expected expression profile. It was grouped into the cambium/xylem cluster in each tissue without significant differences between stems and storage roots. Genes encoding for other important stem cell regulators like *AINTEGUMENTA* (Fig. 4e; *ANT*) or *WUSCHEL-RELATED HOMEOBOX* (*WOX*) *4* (Fig. 4f) were also found in the same expression clusters. ANT directly controls cell proliferation in a cytokinin-dependent manner together with D-type CYCLIN D3;1 (Mizukami and Fischer, 2000; Randall *et al*., 2015). WOX4, on the other hand, acts downstream of PXY to control the position of stem cell organizer of the vascular cambium (Etchells *et al*., 2013; Ji *et al*., 2009). All these cambium markers are important either for its location or capacity to produce new cells. It is therefore expected to see similarities in expression between stems and storage roots.

In contrast, striking differences were found for xylem marker genes involved in the formation of secondary cell walls, which is largely controlled by a gene regulatory network involving different NAC and MYB transcription factors that are expressed in developing xylem cells (Wang *et al*., 2011; Zhong *et al*., 2010; Zhong *et al*., 2008). These include NAC SECONDARY WALL THICKENING PROMOTING FACTOR (NST) 1-3 which are important for fiber formation. MYB46/83 act downstream of NST and VND as master switches for SCW formation. *MeNST1* (Fig. 4g), *MeMYB46* (Fig. 4h), and *MeMYB83* (Fig. 4i) were all significantly lower expressed in the storage root compared to the stem. Still, *MeNST1* and *MeMYB46* could be detected in stem and storage root xylem, but the expression of *MeMYB83* in storage roots did not reach the minimum expression threshold to be considered a DEG (> 50 average counts). The lower expression of secondary cell wall regulators in storage root xylem was expected, as the reduction in secondary cell wall formation is one of the obvious features of storage parenchyma cells. Furthermore, these results show that it is possible to find differentially expressed genes with relevant functions using this analysis approach and that further analysis should reveal more players involved in storage parenchyma formation.

### Expression of genes involved in secondary cell wall biosynthesis differs between tissues

One major aspect of storage root development is the lack of secondary cell walls of xylem cells. To study this in more detail, the analysis of the regulatory network of secondary cell wall formation was expanded to include the expression profiles of additional key transcription factors (Fig. 5, Supplementary File 6) as well as enzymes involved in secondary cell wall and lignin biosynthesis (Fig. S1, Fig. S2, Supplementary File 6).

**Figure 5.**
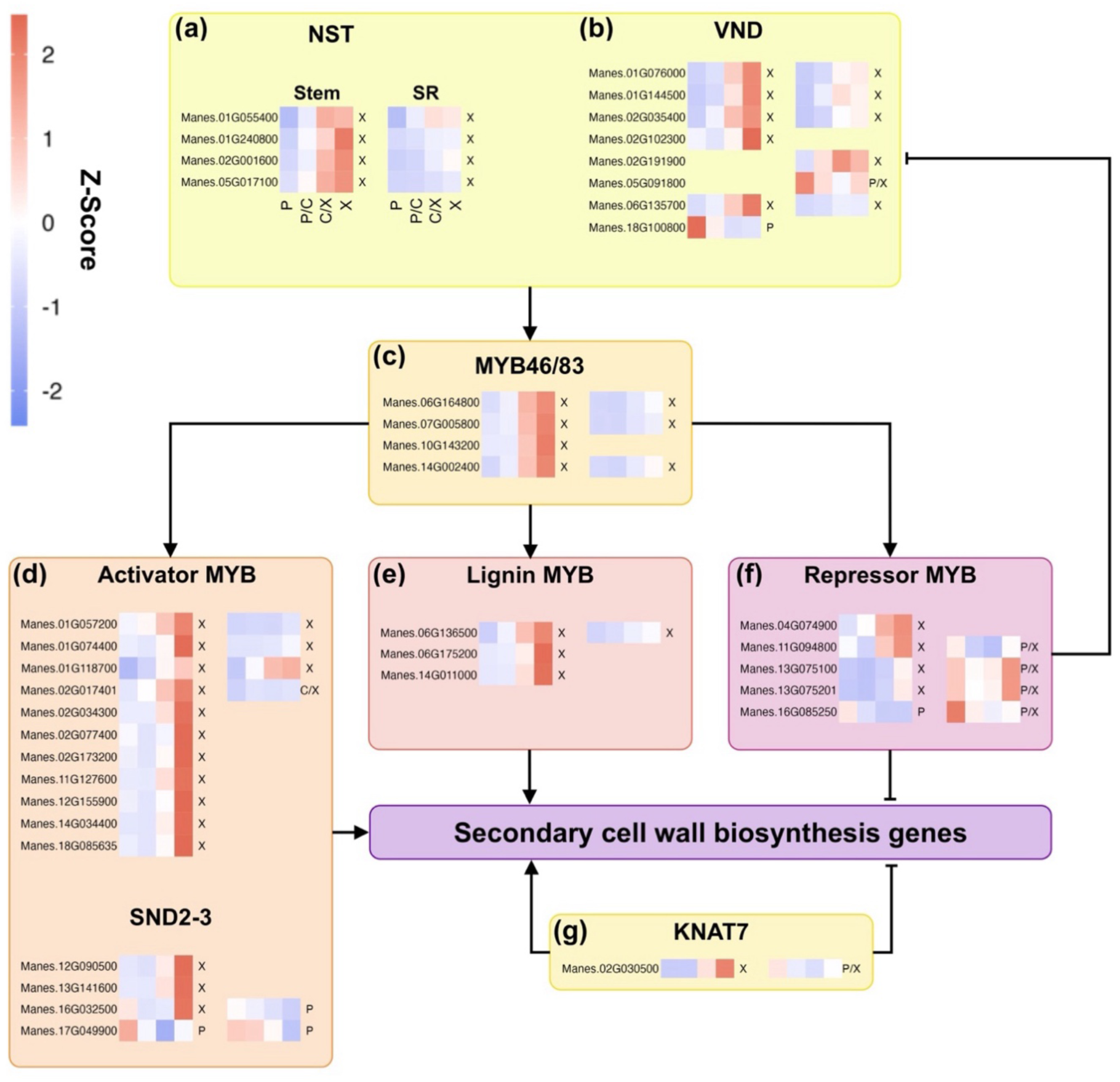
Expression of transcription factors involved in secondary cell wall deposition. Schematic representation of the regulatory network controlling secondary cell wall biosynthesis. Heatmaps display row wise z-scores of each gene encoding for a cassava ortholog of the *A. thaliana* proteins (a) NST1-3, (b) VND1-7, (c) MYB46/83, (d) MYB20/42/43/52/54/69/103 and SND2/3, (e) MYB58/63/85, (f) MYB4/7/32 (g) KNAT/KNOX7. Only genes that were clustered in either tissue are shown. Left and right heatmaps indicate stem and storage root, respectively. Within each heatmap, the tiles from left to right correspond to the sample clusters A (Phloem), B (Phloem/Cambium), C (Cambium/Xylem) and D (Xylem). Letters indicate the cluster. P: phloem, C: cambium, X: xylem.

Xylem fiber and vessel initation is controlled by a distinct group of NAC transcription factors. These include *NST* genes that were identified in *A. thaliana* (*AtNST1-3*) as key regulators specific for fiber formation (Zhong *et al*., 2006; Zhong *et al*., 2010; Zhong *et al*., 2008). Four *MeNST* genes were found, all of which were grouped within the xylem expression cluster of both stem and storage root, albeit exhibiting lower expression levels in the latter (Fig. 5a). This is consistent with the decreased number, but not complete lack of fibers in the storage organ (Fig. 1). Tracheary element initiation in *A. thaliana* is governed by a group of seven VND transcription factors. Eight cassava *VND* genes were identified (Fig. 5b), five of which displayed similar expression profiles to the *MeNST* genes. Notably, one ortholog was undetectable in stems, but exhibited weak expression in storage root xylem, while another was expressed in both storage root phloem and xylem, with one being exclusive to stem phloem. Three out of the four the *MeMYB46*/*83* genes (Fig. 5c), which are master regulators of secondary cell wall formation downstream of the NST/VND cassette were expressed in both stem and storage root xylem. Similar to the *MeVND* genes, all but one were significantly lower expressed in the storage organ, while the last was undetectable in storage roots. Further downstream, a group of MYB and NAC transcription factors comprised of *AtMYB20, 42, 43, 52, 54, 69* and *103,* as well as *AtSND2/3* act as positive regulators of SCW formation (Zhong *et al*., 2008). Eleven cassava orthologs of these MYB activators were DEGs and could all be found in stem xylem and only three in storage root xylem (Fig. 5d). *AtMYB*58, *63,* and *85* transcription factors are involved in lignin biosynthesis specifically. All three cassava ortholog were expressed in stem xylem, but only one could be detected in storage root with expression in the xylem (Fig. 5e).

The majority of positive regulators of secondary cell wall deposition follow the same expression profile as the xylem marker genes (*NST1*, *MYB46*, *MYB83*) with expression in the xylem of stem and storage root tissues, but with clearly higher expression in stem xylem over storage root xylem (Fig. 4g-i). However, orthologs of transcription factors involved in negative regulation of the regulatory network behaved differently. These include orthologs of *AtMYB4, 7,* and *32* as well as KNOTTED-LIKE HOMEOBOX PROTEIN FROM ARABIDOPSIS THALIANA (KNAT) 7 (Fig. 5f-g). While all five *MeMYB* genes in this group were expressed in stem xylem, four of them displayed higher expression in phloem and xylem of the storage root (Fig. 5f). Moreover, three of them were significantly higher expressed in storage root compared to the stem. The only *MeKNAT7* gene exhibited a comparable expression profile between stem and storage root tissue, but was lower expressed in storage roots (Fig. 5g). Genes encoding for enzymes involved in hemicellulose (Fig. S1) and lignin biosynthesis (Fig.S2) were mostly co-expressed with the secondary cell wall regulatory network as expected. However, exceptions were observed among genes involved in multiple pathways, such as the CINNAMYL ALCOHOL DEHYDROGENASE (CAD) genes, which have additional roles beyond monolignol biosynthesis, including glutathione biosynthesis (Fig. S2).

Taken together, these findings suggest that the reduced secondary cell wall formation in the xylem of the storage root is mainly controlled through the reduced expression of the required activators, but also the involvement of a distinct set of differently regulated genes likely suppressing expression of the genes involved in secondary cell wall biosynthesis.

### Storage roots display differences in the expression profiles of key vascular cambium regulators

All secondary xylem and phloem cells are produced in the cambial zone of both stem and storage roots (Fig. 1). Therefore it is likely that differences in the xylem and xylem transition zone, which cause the repression of secondary cell wall production in the storage root are accompanied by differences in the cambial zone as well. However, the initiation and maintencance of the cambium represents a more complicated and far less understood process than secondary cell wall formation and involves a plethora of different transcription factor families and plant hormones (for recent reviews see Fischer *et al*. (2019); Turley and Etchells (2021)). For this reason, only some will be discussed in detail here (an extended list of potentially important genes can be found in Supplementary File 6).

Both stem and storage root cambia share crucial regulatory mechanisms, such as the expression of *MePXY* and *MeWOX4*, which are predominantly expressed in the cambium/xylem samples (Fig. 4d, f). The position of the vascular cambium within the tissue is controlled through a local auxin maxima, which controls the localization of PXY through the activity of HD-ZIPIII transcription factors (Etchells *et al*., 2013; Etchells and Turner, 2010; Smetana *et al*., 2019). All *MeHD-ZIPIII* genes were expressed in the cambium/xylem cluster of both tissues with similar expression levels. The specific auxin signalling component for vascular cambium development include orthologs of *A. thaliana AUXIN RESPONSE FACTORS (ARF) 7*, *19*, and *MONOPTEROS* (*MP*), which also showed the same expression profile in both tissues. This gives the overall impression that the most basic processes of the cambial zone are shared between stem and storage root.

Conversely, notable difference could be observed for genes involved in the differentiation control of the cambial zone (Fig. 6). These include orthologs of the important regulator *BREVIPEDICELLUS/KNAT1* (*KNOX1;* Fig. 6a-b). KNOX1 is a class 1 KNOTTED-LIKE protein that plays an important role in the maintenance of the vascular cambium and differentiation of xylem cells based on its expression level within a cell (Liebsch *et al*., 2014). *KNOX1* genes usually show an expression gradient that is higher on the phloem side of the cambium and decreases towards the xylem (Schrader *et al*., 2004). This is also the case for cassava stems. However, two of three *MeKNOX1* genes showed a different expression profile in the storage root where they were expressed in both phloem and xylem (Fig. 6a-b), which corresponds to the majority of parenchyma cells (Fig. 1), indicating their involvement in storage parenchyma formation. Interestingly, GO term enrichment (Supplementary File 7) showed an enrichment of certain auxin signalling genes within genes higher expressed in the storage root phloem/xylem, which suggests a change in auxin signalling between stem and storage root.

**Figure 6.**
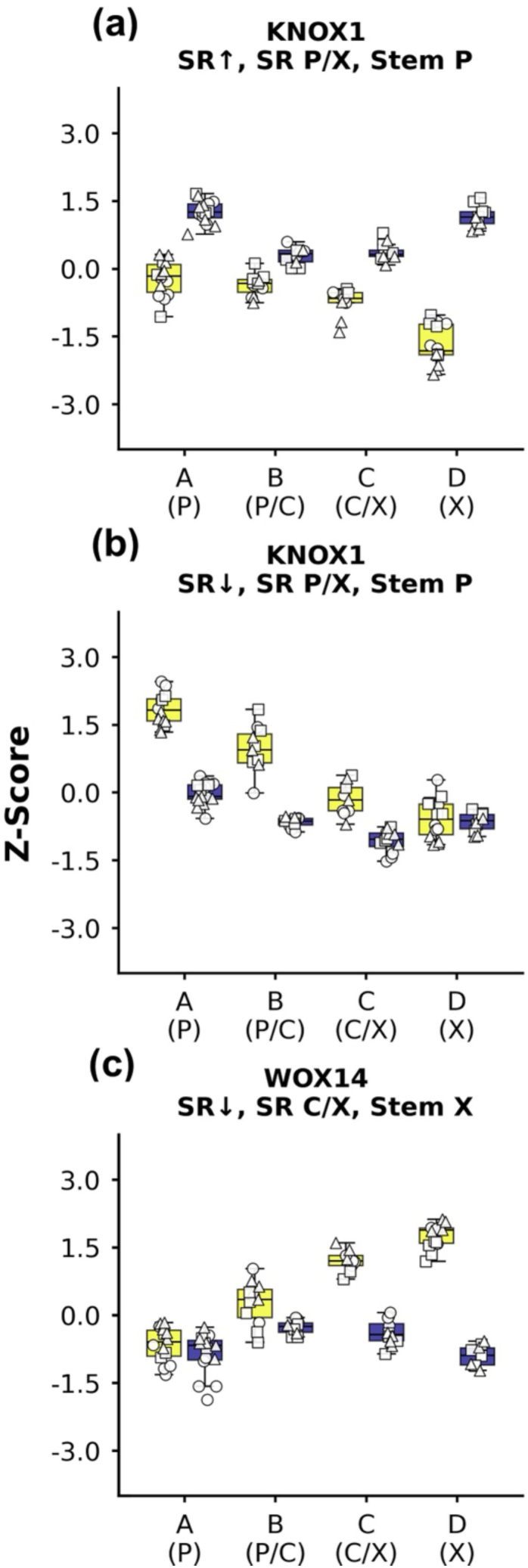
Well-known cambium genes with deviating expression profiles. Shown is the expression of key regulators with notable expression differences between stem and storage root against the sample cluster. Data is centered and scaled based on batch corrected VST values. The displayed genes were manually picked from all putative vascular cambium regulators based on their remarkable expression profile: (a) Manes.05G184900, (b) Manes.18G052901, (c) Manes.05G124300. Panel titles indicate the trivial name and cluster association. Abbreviations: SR: storage root, SR↑: higher in SR, SR↓: lower in SR, P: phloem, C: cambium, X: xylem.

Contrary to *MeKNOX1*, the sole *MeWOX14* gene displayed an increase towards the stem xylem in stems, while being only expressed in the storage root cambium/xylem sections and generally much lower compared to the stem (Fig. 6c). WOX14 plays a role in xylem differentiation through gibberellic acid (GA) signaling (Denis *et al*., 2017). In *A. thaliana* hypocotyls, GA promotes the transition from xylem parenchyma to xylem fiber formation during xylem expansion. Cassava storage roots display very little xylem fiber formation. Therefore, decreased WOX14 expression in storage root xylem might be an important part of storage parenchyma formation. However, GA is not necessarily produced locally and no clear trend could be found for any GA biosynthesis or signalling genes.

### Analysis of interesting expression patterns likely influencing parenchyma formation

The combination of unbiased clustering and the biased analysis of individual genes and pathways allowed to identify potential key players and pathways involved in storage root development of cassava. Using this information, a set of expression profiles was defined to be further analyzed, specifically to find additional transcription factors and pathways outside the traditional cambium players (Supplementary File 8). These include (1) genes co-expressed with secondary cell wall biosynthesis genes that are expressed either only in stem xylem or the xylem of both tissues, but significantly higher in the stem (MYB46; 2,387 DEGs). (2) Genes with an expression profile similar to *MeKNOX1* with higher expression at the phloem side of the vascular cambium and decreasing expression towards the xylem in the stem and a “U-shaped” expression profile in the storage root, matching the pattern of most parenchyma cells in both stems and storage roots (KNOX1; 1,530 DEGs). (3) Genes with the inverse KNOX1 pattern, co-regulated with *MeWOX14* and expressed mainly where few parenchyma cells were produced (WOX14; 1,693 DEGs).

A GO term enrichment analysis (Supplementary File 9) showed that the MYB46 group was strongly enriched in biological processes involved in secondary cell wall biosynthesis and the same terms that were enriched in the individual xylem clusters (i.e. “plant-type secondary cell wall biogenesis”, “positive regulation of secondary cell wall biogenesis”, “lignin biosynthetic process”, etc.). This was expected, but shows the dominance of these pathways within this expression pattern. The KNOX1 group, on the other hand, was dominated by biological processes around different stress responses including “response to water deprivation”, “response to cold” and “response to salt stress” (Supplementary File 10). This is in line with previous findings, where water regulation and osmolyte accumulation in vacuoles was indicated to be an important step in storage parenchyma formation (Rüscher *et al*., 2024). In addition, “post-embryonic plant morphogenesis” consisting of *LIGHT SENSITIVE HYPOCOTYL* (*LSH*) genes that are involved in the development of organ boundaries and part of the same regulatory network as KNOX1 were found with similar expression profiles (Fig. S3), further strengthening the case for the importance of this particular network in parenchyma formation (Cho and Zambryski, 2011).

In a next step we aimed to find potential transcription factors that are involved in the regulation of these expression patterns. As expected, transcription factor binding sites (TFBS) of many of the MYB transcription factors were enriched in the MYB46 group (Fig. 7; Supplementary File 11). Four of those genes were also part of the same expression group and were all part of the MYBs involved in secondary cell wall formation as shown in Figure 5. The KNOX1 group also showed a clear pattern. TFBS of 10 different BASIC LEUCIN ZIPPER (bZIP) transcription factors were enriched and three of those were also expressed in the KNOX1 group (Fig. 7). These include *Manes.04G074300*, an ortholog to *AtbZIP44*, which falls into the SUCROSE INDUCED REDUCTION IN TRANSLATION (SIRT) group (Dröge-Laser *et al*., 2018; Rahmani *et al*., 2009; Sagor *et al*., 2016). Members of this group contain an upstream open reading frame (uORF) important for sucrose-dependent regulation, which was also found in *Manes.04G074300,* as well as orthologs of other *A. thaliana* SIRT bZIP proteins (Fig. S4). The other two enriched and expressed cassava *bZIP* genes are both involved in abscissic acid (ABA) signalling, encoding for a G-BOX BINDING FACTOR 3 (GBF3; Manes.01G249900) and an ABA RESPONSE ELEMENT BINDING FACTOR 1 (ABF1/AREB1; Manes.18G037900) homolog (Fujita *et al*., 2005; Lu *et al*., 1996). Furthermore, multiple other *MebZIP* genes were found in the KNOX1 group including another *GBF3* (Manes.05G026700) and *AREB1* homolog (Manes.05G174500, Manes.18G037900) together with an *ABF3* gene (Manes.10G080700). Alongside the expression and enrichment of TFBS of transcription factors involved in ABA signalling, biological processes that are associated to ABA signalling were enriched in the KNOX1 group (Supplementary file 10). These include responses to different stress condition like salt, cold and water deprivation. The latter specifically comprised around 10 % of all genes in the KNOX1 group and includes important players in the ABA signalling like *SnRK2*, *PP2C* and *LEA* genes (Ng *et al*., 2014). However, unlike the majority of secondary cell wall related genes, which were all co-expressed, other orthologs of the same genes are found with different expression profiles.

**Figure 7.**
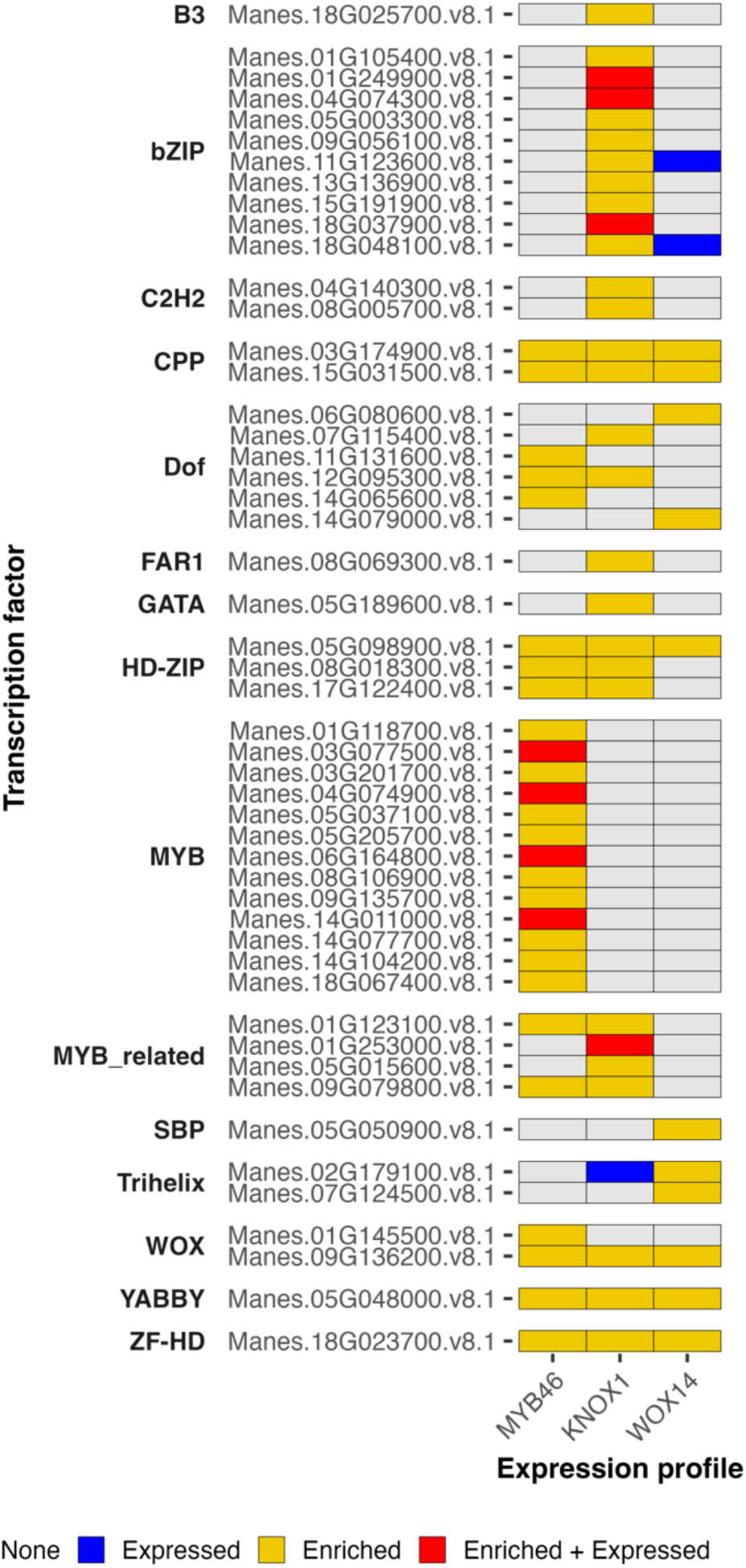
Relevant expression patterns and binding site enrichment. Heatmap depicting transcription factors whose binding sites were enriched in any expression group. Grey indicates neither enrichment nor expression, blue expression but no enrichment, yellow only enrichment but no expression, and red both expression and enrichment in the corresponding tile. X and y axis indicate expression group and Transcription factor respectively. Y-axis headers describe the transcription factor family.

## Discussion

### Cryosectioning successfully captured information about the vascular cambium transcriptome of cassava

Transcriptome analysis of samples harvested through cryosectioning has successfully been used to study the formation of wood development in tree species as it allows for highly-resolved data, while keeping the information about the position within the specimen (Immanen *et al*., 2016; Schrader *et al*., 2004; Sundell *et al*., 2017). This led to a rather good understanding of wood formation and the underlying regulation. However, it was never utilized to study the development of root or tuber storage organs, as the most well-studied storage tissues such as potato tubers do not possess the symmetry needed for this approach. However, cassava storage roots grow from a singular vascular cambium that forms a radial secondary vasculature making them suitable for a cryosection approach. Since storage roots are not straight but convex, the analysis was more difficult compared to stems, as was visible in the higher variance between samples (Fig. 2a). This could be successfully overcome using statistical methods by deducting the batch effects of the individual samples (Fig 2b) and clustering of samples before differential gene expression analysis (Fig. 2c). The clustering and dimensionality reduction did reveal clear separation of samples across the sections (Fig 2d). The gene expression clustering of differentially expressed genes (Fig. 3) and the subsequent analysis of cassava homologs of well-known marker genes from other plants (Fig. 4) demonstrated that the workflow produced reliable expression profiles that reflect the underlying biological process. The presented data represents an important resource for tissue-specific gene expression in cassava stems and storage roots.

### Repression of secondary cell wall biosynthesis is key for storage parenchyma formation

Secondary growth of woody plants is facilitated by the vascular cambium, which forms a roughly cylindrical structure capable of anti- and periclinal cell division that allows trunks and roots to increase in lateral size by depositing secondary phloem outwards and xylem tissue inwards (Fischer *et al*., 2019; Nieminen *et al*., 2015; Turley and Etchells, 2021). The anatomy of the vascular cambium, however, is more complex and consists of multiple different zones and cell types (Lachaud *et al*., 1999; Nieminen *et al*., 2015). Unlike the storage root that is mainly comprised of storage parenchyma, cassava stems produce large amounts of wood consisting of water conducting tracheary elements, stabilizing fibers and parenchyma cells (Rüscher *et al*., 2024), typical for any tree or shrub (Furze *et al*., 2018; Schuetz *et al*., 2012). These cells were produced in the xylem transition zone (Fig. 1b) where cell division stopped, tracheary element precursors rapidly inflated, and cell walls of fibers started thickening. However, no lignin was deposited yet visualized by the lack of the indicative blue staining of toluidine blue treated samples (Fig. 1b). The transition zone of the storage root also contained developing tracheary elements, but lacked the majority of developing fibers (Fig. 1a). Instead, these cells began swelling and to store starch after the end of the division, requiring some differential regulation of secondary cell wall biosynthesis (Fig. 1a).

Fittingly, these differences in morphology were accompanied by clear disparities in expression between the two tissues. Transcripts of the master- and positive regulators of secondary cell wall biosynthesis (Wang *et al*., 2011; Zhong *et al*., 2017, 2019; Zhong *et al*., 2006; Zhong *et al*., 2010; Zhong *et al*., 2008) shared the same expression profile by being higher expressed towards the xylem of both stem and storage root tissues, but overall less abundant in the storage root (Fig. 4g-i, 5). The same trend was found for genes encoding for enzymes involved in (hemi-)cellulose and lignin biosynthesis (Fig. S2-3). Nevertheless, most of the secondary cell wall related genes were still detectable in the storage root, probably reflecting the reduced quantity, but not the absence of lignified cells. Furthermore, implying that positive regulation of secondary cell wall biosynthesis works similarly across tissues. Besides quantitative differences, qualitative differences could be found in putative negative regulators of secondary cell wall formation. These were both expressed in storage root phloem and xylem, while being expressed only in the xylem of stems (Fig. 5f). Furthermore, they showed higher expression in the storage organ, fitting to the reduced number of lignified cells.

Overall, the repression and differential regulation of genes involved in secondary cell wall biosynthesis is a key aspect of storage parenchyma formation, which is reflected both transcriptionally and morphologically. Nevertheless, the observed changes occur high up in the regulatory network’s hierarchy, which makes changes earlier during differentiation in the regulation of the cambial zone likely.

### Key cambium regulators and phytohormones control aspects of storage parenchyma formation

The cambial zone consists of two different cell types called ray and fusiform initials that produce xylem rays and the actual phloem/xylem cells, respectively (Lachaud *et al*., 1999; Nieminen *et al*., 2015). Both were found in the cambial zone of cassava stems, as well as storage roots making it rather similar between tissues (Fig. 1). This similarity was further supported by a comparable transcriptome of the cambium/xylem clusters of stems and storage roots (Fig. 3). However, a large overlap of genes significantly higher expressed in storage roots and found in both cambium/xylem expression clusters of stems and storage roots could generally be annotated as part of cell cycle or cell division functions (Supplementary File 5), likely reflecting the increased cell proliferation of the storage root in comparison to the stem.

Genes involved in vascular cambium formation and maintenance were expressed similarly between stem and storage roots, including, but not limited to *WOX4, ANT* and, *PXY* genes (Fig. 4). The position of the vascular cambium is defined by a local auxin maximum that defines the position of the stem cell organizer, which expresses the *HD-ZIPIIIs* and *PXY* (Etchells and Turner, 2010; Smetana *et al*., 2019). The ARFs involved in cambium maintenance are *MP*, *ARF7* and *ARF19* whose corresponding cassava orthologs were co-expressed with *PXY* and showed no differences between tissues. Nevertheless, there was evidence for a difference in auxin signaling between stem and storage roots, because of a GO term enrichment of “response to auxin” specifically in genes expressed higher expressed in storage roots with expression in both phloem and xylem (Supplementary File 7). This pattern describes the storage parenchyma and was shared by *MeKNOX1* genes (Fig. 6). These already gained attention in previous studies for their remarkable expression profile compared to other vascular cambium maintenance genes by showing an increase in expression upon root bulking (Rüscher *et al*., 2021; Sojikul *et al*., 2015). Unlike in the storage root, *MeKNOX1* expression was strongest in the phloem and decreased towards the xylem in stems (Fig. 6a-b). This is in line with the described expression profile in other cryosectioning data of aspen trunks (Schrader *et al*., 2004) and congruent with reporter gene assay data from *A. thaliana*, which describe *AtKNAT1* as being expressed mainly in the cambium with a gradient towards the xylem. This gradient appears to be important to prevent cambial cells from differentiating into lignified cells and developing xylem cells from further division (Liebsch *et al*., 2014). This model could explain the *MeKNOX1* expression in storage roots (Fig. 6a-b), where this gradient towards the xylem does not exist, but expression increases again towards the xylem, therefore preventing differentiation into sclerenchyma. Furthermore, *MeKNOX1* generally follows the storage parenchyma in the storage root, which is produced towards the xylem and phloem. Other notable genes in the KNOX1 pattern include most *LIGHT-SENSITIVE HYPOCOTYL* (*LSH*) genes. *Arabiodpsis LSH* proteins are important for organogenesis and boundary formation and are part of the gene regulatory network involving *AtBP/KNAT1,* further implying a prominent role of the *MeKNOX1* regulatory network in storage parenchyma formation (Cho and Zambryski, 2011).

Conversely to the *KNOX1* expression pattern, the sole *MeWOX14* gene was expressed higher towards the xylem in the stem, but decreased towards the phloem. In storage roots, no such gradient was observed and expression was generally much lower expressed in the storage root (Fig. 6c). Just like WOX4, WOX14 is thought to act downstream of PXY, but does not appear to be directly involved in the formation and maintenance of the meristematic stem cells (Etchells *et al*., 2013). In Arabidopsis, *wox14* mutants demonstrate a milder phenotype compared to *wox4* mutants, characterized by delayed flowering, whereas overexpression results in increased production of lignified xylem cells (Denis *et al*., 2017). The expression of *MeWOX14* in stem xylem implies a similar role in cassava wood formation. Conversly, reduction of *MeWOX14* expression in storage root xylem is likely an important aspect of storage parenchyma formation. Notably, the late flowering phenotype of *wox14* mutants can be rescued by exogenous application of bioactive GA (Denis *et al*., 2017). GA is involved in the shift from parenchyma to fiber formation during xylem expansion and is most abundant in the xylem side of the vascular cambium (Björklund *et al*., 2007; Immanen *et al*., 2016; Ragni *et al*., 2011). However, the exact region of biosynthesis is rather unclear as the expression domain of the corresponding biosynthesis genes stretches from phloem to expanding xylem (Israelsson *et al*., 2005). Furthermore, GA also works as a mobile signal to signal the transition towards xylem expansion (Björklund *et al*., 2007; Ragni *et al*., 2011). It is therefore not surprising that no clear trend was observed in GA biosynthesis or signaling components in the here presented data. However, reduction on a whole root level throughout bulking was observed multiple times (Rüscher *et al*., 2021; Sojikul *et al*., 2015). Overall, due the lack of fiber formation and *MeWOX14* expression in the storage root xylem, the repression of GA signaling is likely a key part of storage parenchyma formation.

While GA seems to inhibit storage organ formation, ABA was shown to promote storage parenchyma formation in potato tubers (Xu *et al*., 1998). In the cassava storage root parenchyma, ABA signaling components and AREB TFBS were found to be enriched. An earlier study demonstrated extensive regulation of vacuolar processes in cassava storage root bulking, likely due to the vacuoles involvement in cellular growth processes (Rüscher *et al*., 2024). Vacuolar-related processes were reflected in the enrichment of GO terms related to abiotic stresses (Supplementary File 10) and are generally controlled through ABA (Fujita *et al*., 2005; Ng *et al*., 2014). Therefore, the ABA-related signaling might be mostly connect to the regulation of cellular expansion in parenchyma cells (Fig. 8).

**Figure 8.**
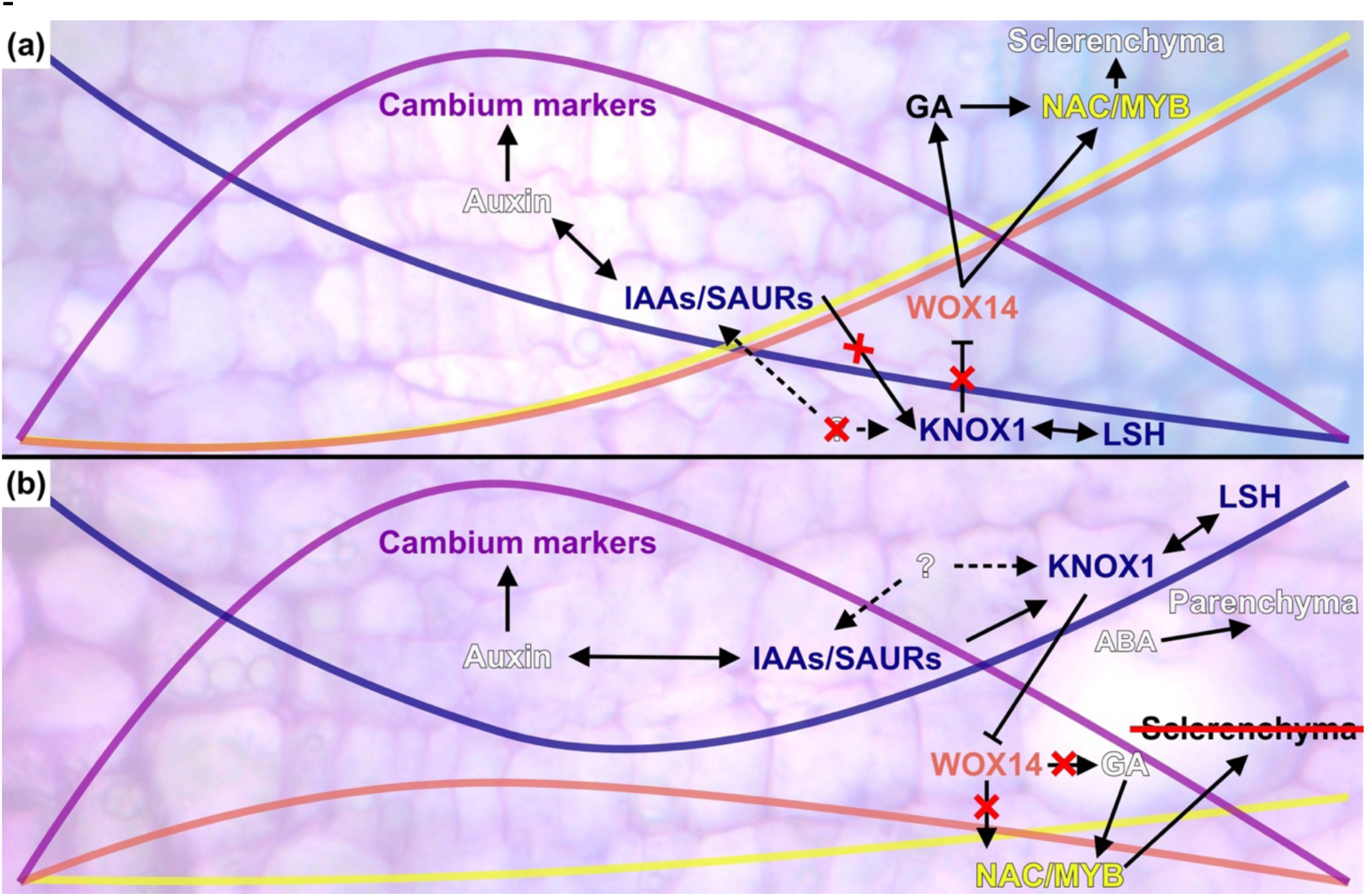
Proposed mechanism of storage parenchyma and wood formation in cassava. Schematic depiction of the formation of (a) wood in the stem, and (b) storage parenchyma in the storage root of cassava. Colors indicate different expression profiles: blue = KNOX1, yellow = MYB46, red = WOX14, purple = cambium markers. Micrographs are the same as in Figure 1.

## Conclusion

Storage parenchyma formation is complex and controlled by many as of yet unknown factors. Our data suggests that key factors of cassava storage parenchyma formation are the repression of GA-signaling and the NAC/MYB regulatory network that is involved in secondary cell wall deposition, as well as changes in the expression profile of important differentiation regulators like KNOX1 and WOX14. Additionally, specific auxin and ABA signaling components in the storage root xylem parenchyma might contribute to the increased cell division and cell expansion observed for the storage organ. Our current understanding of storage root parenchyma formation is summarized as a model in Figure 8.

## Material and methods

### Plant material and cryosectioning

Cassava plants (genotype TME7) were planted in 14 l pots from 20 cm long stakes in a glasshouse in Erlangen, Germany. Fourteen plants were harvested after two months. Whole developing storage roots and the base of the largest newly sprouted stem were wrapped into foil and immediately frozen in liquid nitrogen. For cryosectioning, approx. 0.5 cm long pieces of a single stem internode or a roughly straight storage root piece were prepared from the whole specimen. The pieces were trimmed by hand to around 1-2 mm at the site of sectioning. A Leica CM 3050 S cryomicrotome was used for sectioning at −25 °C. Longitudinal sections were taken until the blade was close to the vascular cambium. Subsequently, 25x 20 µm thin sections were taken, put into 50 µl RNase-free water in a 2 ml reaction tube and immediately frozen in liquid nitrogen. Cross sections of the top of the sectioned piece were taken before and after sampling, stained for a few seconds in 0.1 % toluidine blue and pictures taken under a light microscope. 20 µm wide sectors were added to the before-sectioning images starting at the part closest to the microtome blade and compared to the after-sectioning images. Only, when the start of the after-sectioning and the end of the last sector of the before-sectioning image aligned and the sections reached deep enough into the xylem, where the samples deemed fit for further analysis, resulting in a total of 7 applicable stems and storage roots. Of those three were chosen randomly. We aimed for a span from −100 µm to +220 µm away from the center of the cambial zone. However, due to the nature of sampling this was not possible for all resulting in a total of 95 samples.

### RNA Sequencing and raw data processing

RNA was extracted using the RNeasy Mini Kit (Qiagen). RNA was sent to a service provider for low-input paired-end mRNA sequencing (> 20 million reads, PE150). Resulting FastQ files were quality controlled using FastQC v0.11.9 (http://www.bioinformatics.babraham.ac.uk/projects/fastqc/) and MultiQC v1.13 (https://multiqc.info/). The reads were trimmed for adapter content and quality via bbduk v38.97) (http://sourceforge.net/projects/bbmap/). Bases under a quality of 30 were trimmed from both sides. Reads shorter than 35 base pairs or with an average quality below 30 after trimming were removed. The trimmed reads were mapped to the cassava genome AM560-2 v8.1. (including plastid and mitochondrial genome) with STAR v2.7.10a (Dobin *et al*., 2013). Mapped reads were counted using FeatureCounts v2.0.3 (Liao *et al*., 2014). Only uniquely mapped reads were counted. Multi-overlapping reads were not counted.

### Data analysis

Counts were normalized and log transformed (variance stabilization transformation; VST) using the R package DESeq2 v1.40.1 (Love *et al*., 2014). Subsequent analyses were conducted on each tissue individually. The VST values were batch corrected using LIMMA v3.54.1 and the LRT executed with the plant included as covariate. Downstream analyses were performed on the scaled and centered VST values. UMAP projection was performed in Python using the umap-learn module v0.5.3. Samples were clustered through Louvain community detection with resolution of 1.5 (Blondel *et al*., 2008) in python using the NetworkX module v3.0. For graph generation PCA was performed beforehand on the 5,000 expressed genes with the highest variance (Supplementary file 12). Only genes with a normalized read-depth of more than 50 were considered. The principal components that explain more than 90 % of the total variance – 28 and 32 for stem and storage roots, respectively (Supplementary file 13) - were used to generate a *k*-nearest neighbor graph based on cosine distance (*k* = 7). PCA and the *k*-nearest neighbor were done using scikit-learn v1.02 in python. Clustering results can be found in Supplementary file 14.

The DEG analysis was performed for each tissue individually utilizing an LRT in DESeq2 using the sample cluster and plant as design variables and only plant as reduced model. Genes with FDR < 0.001 were accepted as DEGs. DEGs were also clustered through Louvain community detection with resolution of 1 (stem) and 1.1 (storage root) on a co-regulation network: Edges were drawn between genes based on a Pearson correlation coefficient above 0.8. Genes < 25 edges were removed. Enrichment analysis was performed using a Fisher’s exact test implemented in the R package clusterProfiler 4.8.1. A full list of GO enrichments can be found in Supplementary File 15. If not stated otherwise, plots were generated in R or Python using ggplot2 or matplotlib, respectively. Figures were prepared in Sketch for macOS.

### Functional annotation

Functional annotation data stems from Rüscher *et al*. (2024). Peptides from the *M. esculenta* genome v8.1 were locally blasted (blast+ v2.13.0) against the *A. thaliana* proteome from the araport11 genome (e < 0.001). A list of aliases was downloaded from TAIR (arabidopsis.org, accessed on April 7^th^, 2021). *A. thaliana* genes with potential relevance for cambium development were curated manually from multiple reviews.

### Transcription factor binding site detection

A list of putative cassava transcription factors and binding sites was downloaded from the Plant Transcription Factor Database (planttfdb.gao-lab.org, v5.0; Supplementary File 16) on March 4^th^ 2024 (Tian *et al*., 2019). Promoters of all annotated cassava genes were scanned for those binding sites using FIMO v5.5.5. Promoters were defined as 1000 bp long fragments upstream of the transcription start site indicated in the gff3 file.

## Acknowledgments

We thank Dr. Juha Immanen, Dr. Kaisa Nieminen, Sampo Muranen, and Prof. Dr. Yrjo Helariutta for helpful discussions and their support with the cryosectioning of cassava stems and storage roots.

## Code and data availability

All used commands and scripts are available on github (https://github.com/Division-of-Biochemistry-Publications/CASS/).

Raw sequencing reads have been deposited to NCBI’s Sequence Read Archive (submission ID: SUB14399691) under BioProject ID PRJNA1109716, available at https://www.ncbi.nlm.nih.gov/bioproject/PRJNA1109716.

## Abbreviations

(SCW): Secondary cell wall
(SR): Storage root
(VC): vascular cambium
(XT): xylem transition zone
(RNA-seq): mRNA-sequencing
(PCA): principal component analysis
(PC): principal component
(LRT): likelihood ratio test
(DEG): differentially expressed gene
(VST): variance stabilization transformation
(TFBS): transcription factor binding site

## Author contributions

D.R. performed the greenhouse trials, cryosectioning and microscopy and evaluated the transcriptome data; and did the statistical analysis and visualization of the data. D.R. and W.Z. planned the experiments. U.S. and W.Z. supervised and coordinated the work. D.R. and W.Z. wrote the manuscript with the help of all authors.

