## Supplementary material for "High-resolution spatial transcriptomics of stem and storage root vascular cambia highlights key regulatory processes for xylem parenchyma differentiation in cassava": Ruescher_bioarchives_MS_SupportingInformation.pdf

Running title: Regulation of cassava stem and storage root xylem formation

David Rüscher[1], Uwe Sonnewald[1], Wolfgang Zierer[1*]

[1] Friedrich-Alexander-University Erlangen-Nuremberg, Department of Biology, Division of Biochemistry, Staudtstrasse 5, 91058 Erlangen, Germany

### Supporting Information

#### Supplementary Files

SupplementaryFile1: CSV file containing the results of the LRT test.

SupplementaryFile2: CSV file containing gene clustering results and UMAP.

SupplementaryFile3: CSV file containing the results of the WALD test.

SupplementaryFile4: Excel file containing GO term enrichment results for the intersection of SR Low, Stem X, and SR X.

SupplementaryFile5: Excel file containing GO term enrichment results for the intersection of SR High.

SupplementaryFile6: CSV file of all relevant cassava genes with their corresponding expression profile.

SupplementaryFile7: Excel file containing GO term enrichment results for the intersection of SR High and SR PX.

SupplementaryFile8: Excel file containing all clustered cassava genes with pattern information and expression levels.

SupplementaryFile9: Excel file containing GO term enrichment results for the MYB46 pattern.

SupplementaryFile10: Excel file containing GO term enrichment results for the KNOX1 pattern.

SupplementaryFile11: Excel file containing TFBS enrichment results.

SupplementaryFile10: CSV file containing all putative transcription factors and their expression profile

SupplementaryFile12: CSV file containing PCA used for clustering.

SupplementaryFile13: CSV file containing information about the explained variance of the PCA.

SupplementaryFile14: CSV file containing UMAP and clustering results.

SupplementaryFile15: Excel file containing all GO term enrichment results.

SupplementaryFile16: CSV file containing expression clusters of all cassava transcription factors.

### Supplementary Figures

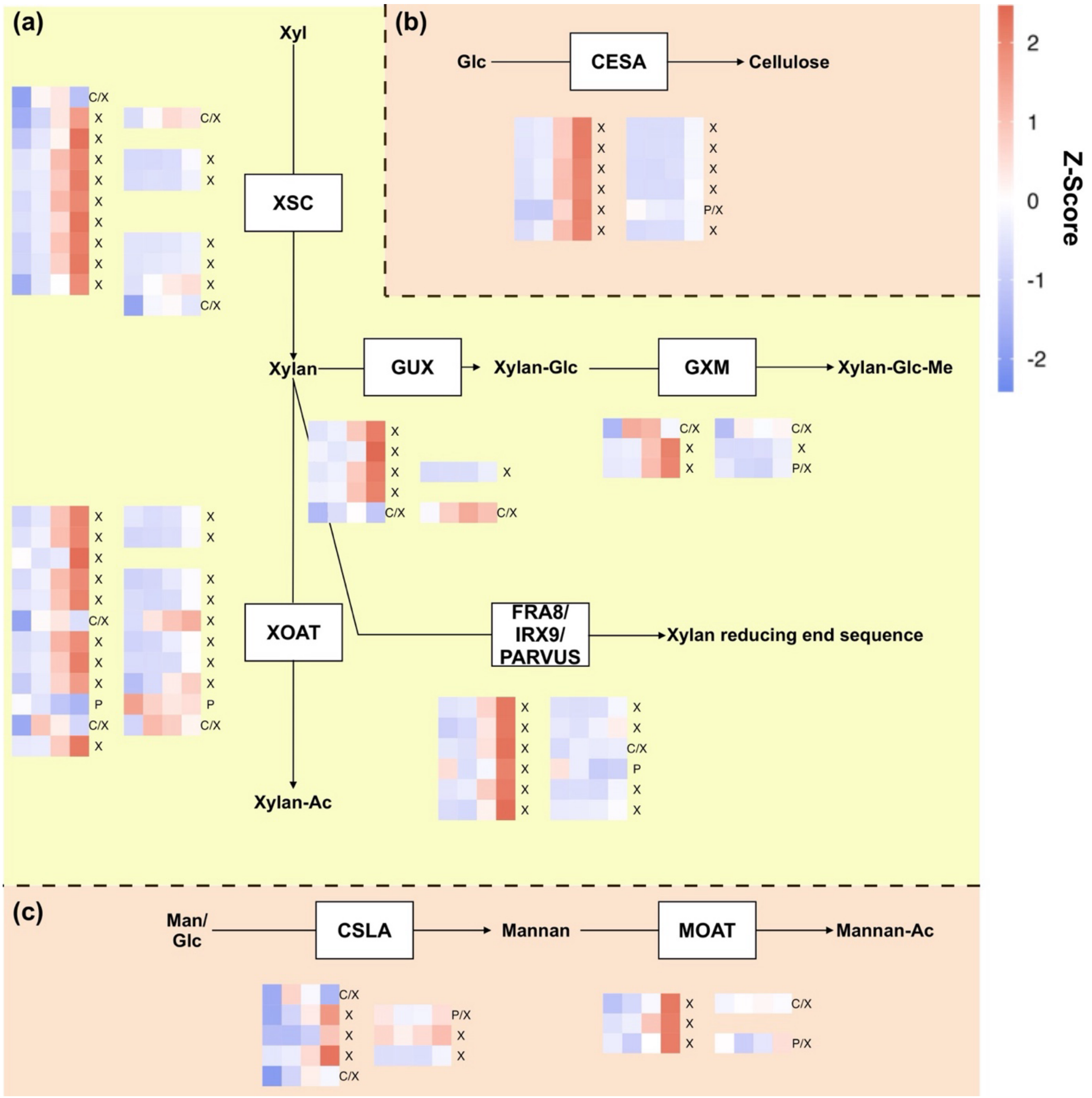


**Fig S1. Expression of genes transcribing for enzymes involved in (Hemi-) cellulose biosynthesis.**
P = Phloem cluster, P/C = Phloem/Cambium cluster, C/X = Cambium/Xylem cluster, X = Xylem cluster. Left boxes represent expression in stems, right boxes represent expression in storage roots.

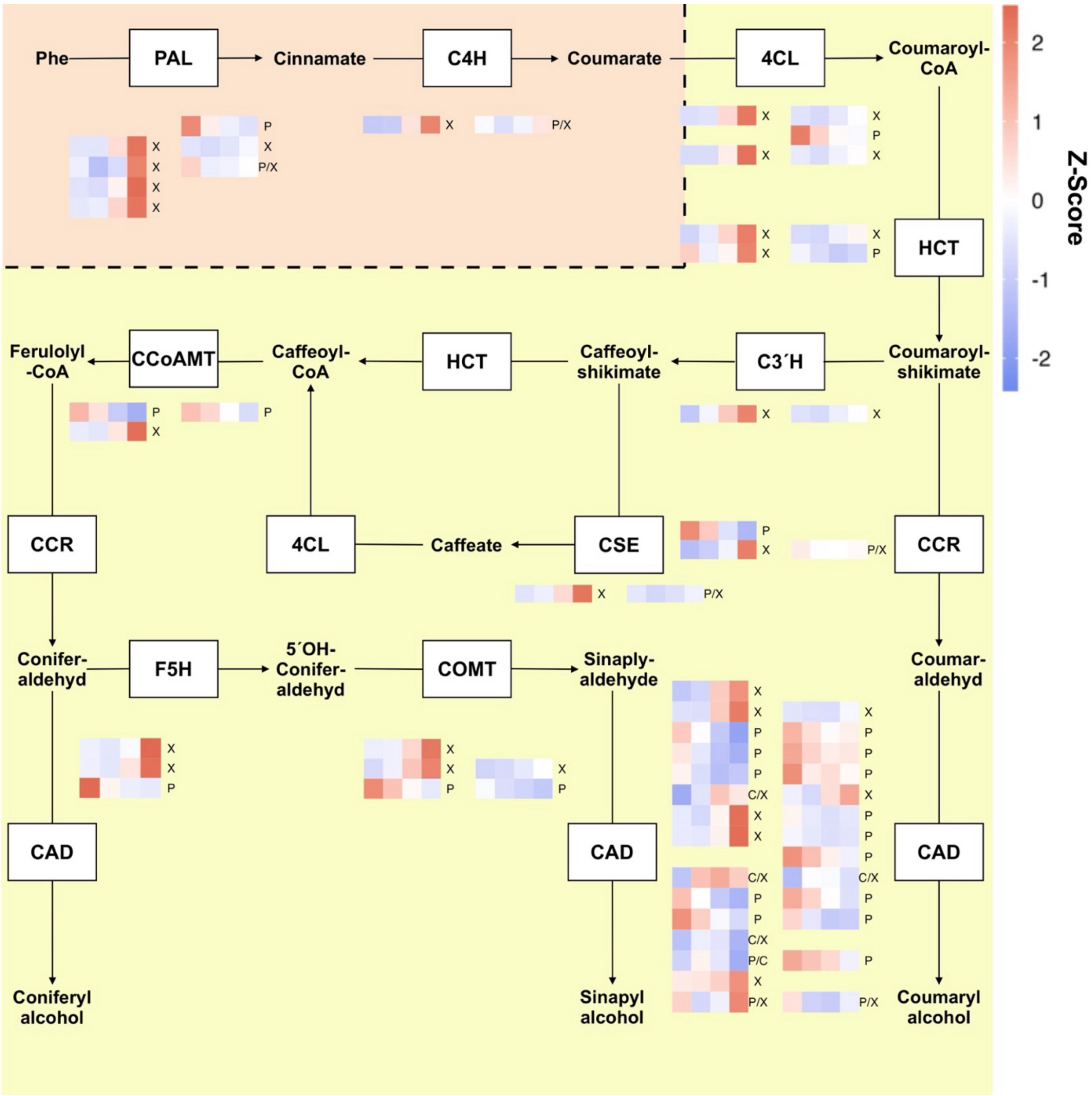


**Fig S2. Expression of genes transcribing for enzymes involved in lignin biosynthesis.**

P = Phloem cluster, P/C = Phloem/Cambium cluster, C/X = Cambium/Xylem cluster, X = Xylem cluster. Left boxes represent expression in stems, right boxes represent expression in storage roots.

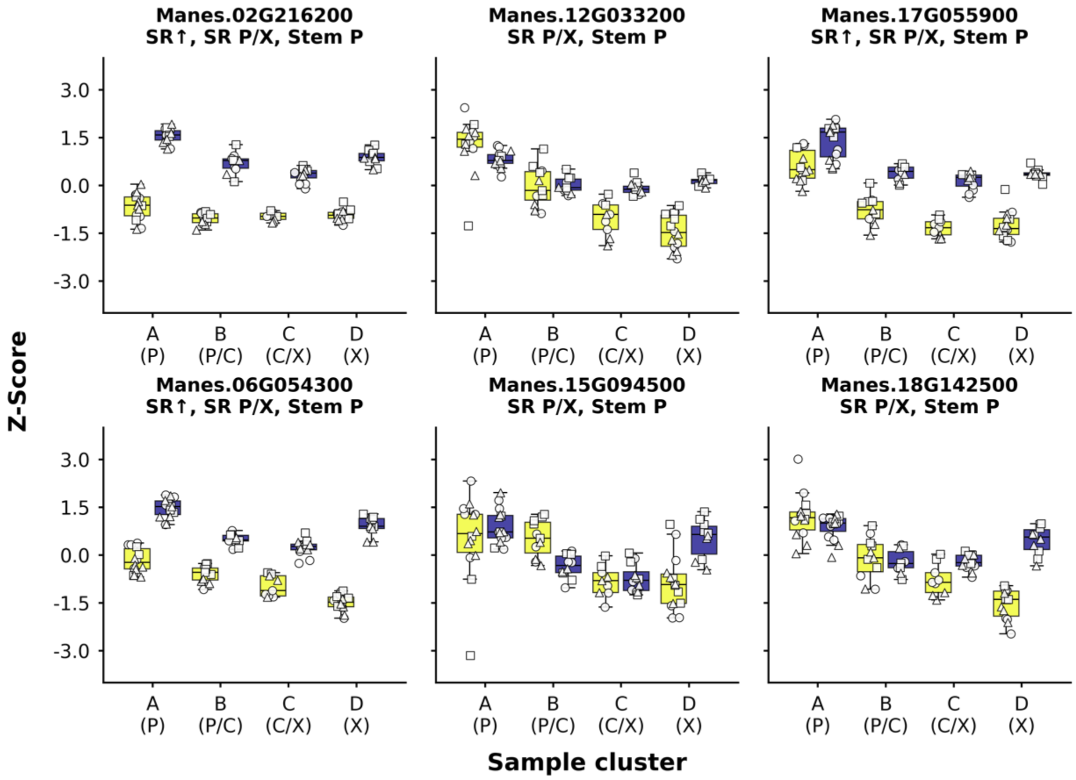


**Fig S3. Expression of *LSH* genes in stems and storage roots.**

P = Phloem cluster, P/C = Phloem/Cambium cluster, C/X = Cambium/Xylem cluster, X = Xylem cluster. Yellow boxes represent expression in stem tissue, blue boxes represent expression in storage root tissue.

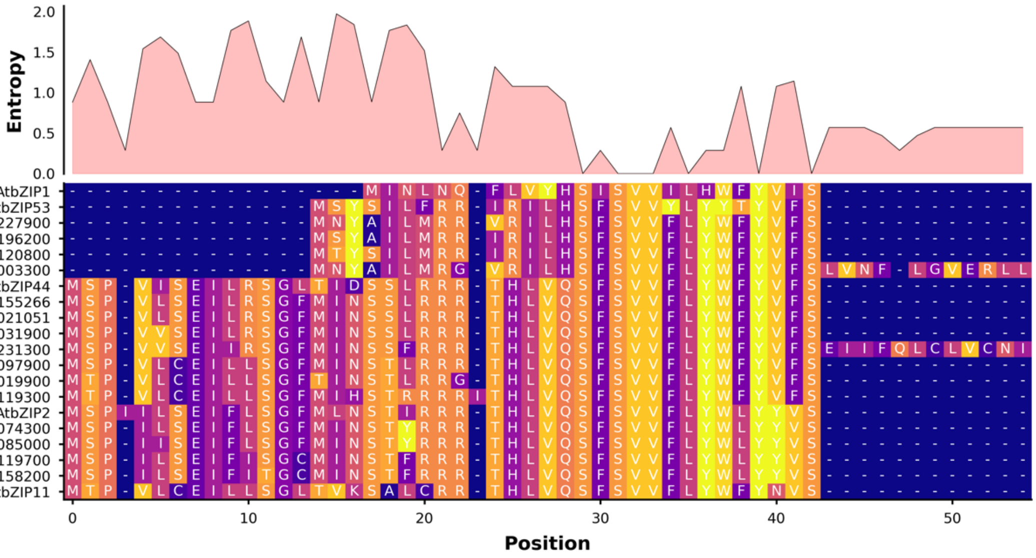


**Fig. S4. Alignment of translated SIRT-bZIP uORF from cassava, poplar, and *A. thaliana.***
