## Supplementary figures and images for "High-resolution spatial transcriptomics of stem and storage root vascular cambia highlights key regulatory processes for xylem parenchyma differentiation in cassava"

### S4_uORF_ALIGNMENT.png

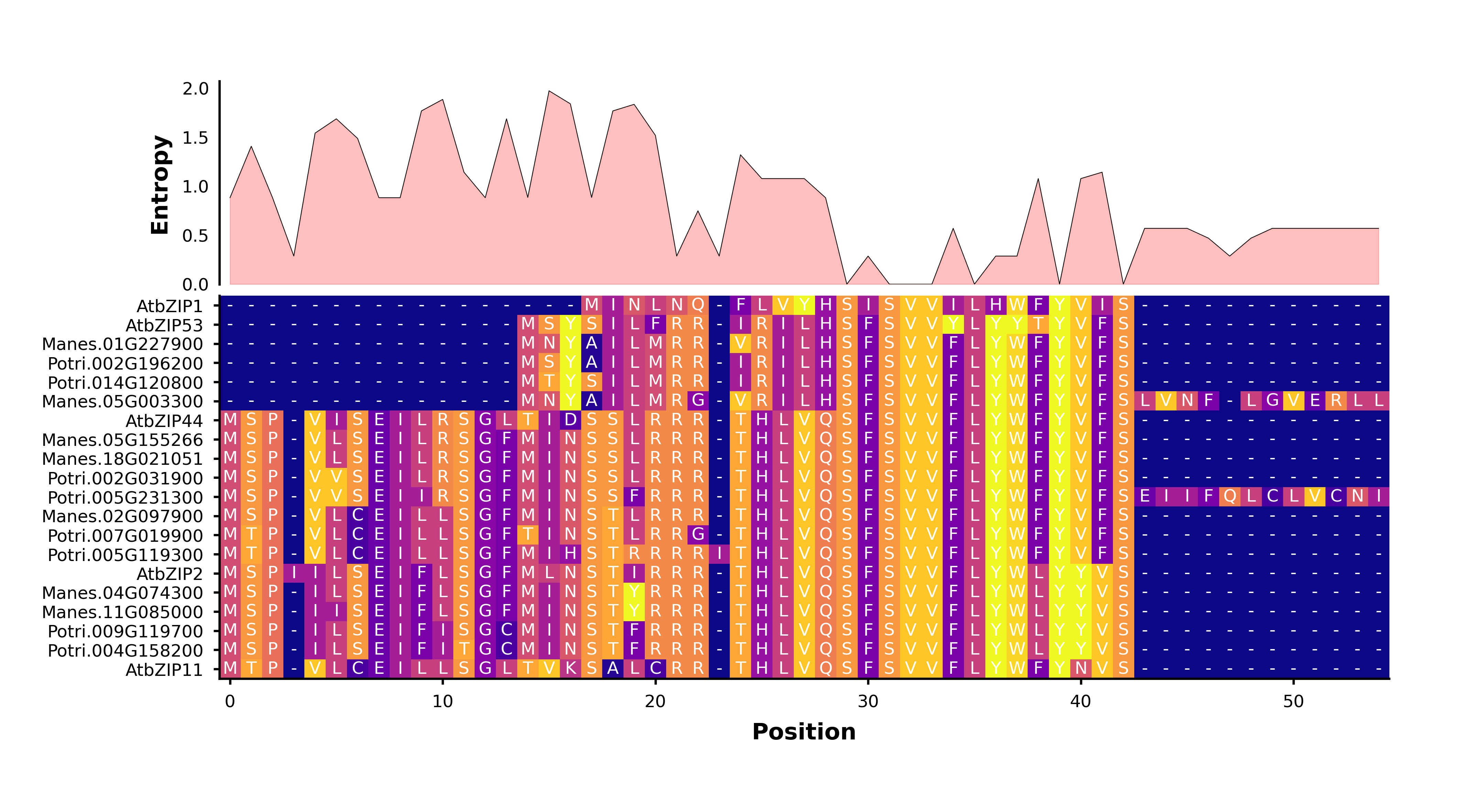
